# The BK channel-NS1619 agonist complex reveals molecular insights on allosteric activation gating

**DOI:** 10.1101/2025.03.27.645783

**Authors:** Naileth Gonzalez-Sanabria, Gustavo F. Contreras, Maximiliano Rojas, Yorley Duarte, Fernando D. Gonzalez-Nilo, Eduardo Perozo, Ramon Latorre

## Abstract

BK channels play essential roles in a wealth of physiological functions, including regulating smooth muscle tone and neurotransmitter release. Its dysfunction, often caused by loss-of-function mutations, can lead to severe phenotypes, including ataxia and sensory impairment. Despite the therapeutic potential of BK channel agonists, the molecular mechanisms by which they stabilize the pore’s open conformation remain unclear. Using cryo-electron microscopy and molecular dynamic simulations, we identified that NS1619, a synthetic benzimidazolone agonist, first described as a BK opener, binds within a pocket formed by the S6/RCK1 linker and the S4 transmembrane segment. Agonist binding drives a twisting motion in the S6 segment, enabling critical interactions with residues K330, K331, and F223. Our findings clarify the mechanism of NS1619 and suggest that its binding site can accommodate other agonists, highlighting a promising target for therapeutic development.

**Teaser:** BK channel activation by NS1619 reveals key binding interactions, offering insights for designing targeted therapeutic agents.

## Introduction

By virtue of its dual activation by intracellular Ca^2+^ and depolarizing voltages, the voltage- and Ca^2+^-activated channel (BK) is the ultimate damper of excitatory stimuli. These characteristics make them essential in numerous physiological functions. In the nervous system, BK channels shorten the action potential, BK by colocalizing with voltage-dependent Ca^2+^ channels, damp neurotransmitter release, and their association with β2 or β4 subunits regulates neuron excitability (*1*, *2*). In smooth muscle, Ca^2+^ sparks produced by the liberation of Ca^2+^ from the sarcoplasmic reticulum promote the activation of BK channels, facilitating muscle relaxation. Hence, they regulate the arterial smooth muscle tone (*3–5*). In the kidney, the BK channeĺs large single-channel conductance is essential to sustain high rates of urinary K^+^ secretion (*6–8*). Not surprisingly, malfunctioning of BK channels in these tissues due to gain- or loss-of-function (GOF/LOF) mutations leads to a variety of pathophysiological phenotypes.

BK channels have been implicated in epilepsy, fragile X syndrome, mental retardation, autism, and chronic pain (*1*, *9*). Analysis of 37 KCNMA1 alleles (*10*) shows that the distribution between patients with GOG and LOF mutations shows the same tendency to seizures. However, paroxysmal non-kinesigenic dyskinesia is predominantly observed among patients with GOF alleles of the BK channel. Other movement disorders are observed in patients with LOF variants. Dysfunction of BK channels related to the loss of the β1 subunit in smooth muscle leads to hypertension (*3*, *4*), an overactive bladder (*11*, *12*), and erectile dysfunction (*13*, *14*). Deleting the pore-forming BK α subunit in the kidney produces an absence of flow-induced K^+^ secretion and hypertension (*7*). Deletion of β1 leads to poor handling of K^+^ loads and is linked with a rise in plasma K^+^ and hypertension. Deleting β4, on the other hand, results in insufficient K^+^ handling and high plasma K^+^, but only mild hypertension (*7*).

Given the large number of LOF BK mutations, identifying BK channel activators with potential therapeutic applications is an important pharmacological target. The first characterized BK channel opener was the synthetic benzimidazolone NS1619 (*15*) (*21*), a potential therapeutic agent in inflammatory pain (*16*), pulmonary hypertension (*17*), erectile dysfunction (*13*), bladder instability (*18*), and migraines (*19*). NS1619 acts as an antagonist in this mutant, rescuing BKG354S cells, a mutation found in a child with congenital and progressive cerebellar ataxia and located in the selectivity filter (*20*). NS1619 appears to activate BK channels via the S6/ RCK1 linker (*22*). However, elucidating the nature and location of the agonist binding site is of fundamental importance in the quest to define its mechanism of action and design drugs able to activate BK selectively and with high affinity. Here, we have determined the structure of the complex BK channel-NS1619 by single particle cryoEM. NS1619 resides in a pocket lined by amino acids from the S6/RCK1 linker and the S4 transmembrane segment. We show evidence suggesting that his arrangement allosterically stabilizes BK’s open conformation by increasing the equilibrium constant that defines the closed-open equilibrium. Furthermore, docking and molecular simulations demonstrate that the NS1619 binding pocket behaves as a robust druggable site, able to accommodate other agonists of a variety of different structures.

## Results

### Calcium-bound Structure Reveal the Putative NS1619-Binding Site

We determined the structure of the human BK channel in the presence of the NS1619 activator using cryo-electron microscopy. Since the effect of the activator is independent of the saturating concentration of Ca^2+^ (*22,* Fig. S1), we followed established purification protocols to obtain the human BK channel in the open state (*23*), adding NS1619 to the sample before freezing under saturating conditions (1 mM final). The structure was solved at a global resolution of 3.4 Å, which allowed detailed modeling of the BK-NS1619 complex. Figure S2 shows the diagram of the global processing and refinement strategy. An additional Coulomb density was observed and attributed to NS1619 (Fig. 1a; red density); this density is absent in previous BK structures observed (23). As anticipated from the functional results, the overall structure of the BK-NS1619 complex remains unchanged when compared with its Apo form, suggesting that NS1619 acts as an open-state stabilizer and confirms the existence of a binding site for NS1619 between RCK1 and the C-linker, which can interact with amino acids from the VSD, in line with molecular docking results. We have identified a set of amino acids that putatively belong to the binding site on the BK channel (all within a 5 Å): Y332, G334, K392, and E399 (Fig. 1 b-d). As shown below, molecular dynamic simulations indicate that this picture changes with time, implying a functional reorganization of the binding site that now include amino acid residues K330, K331, and F223.

**Fig. 1.**
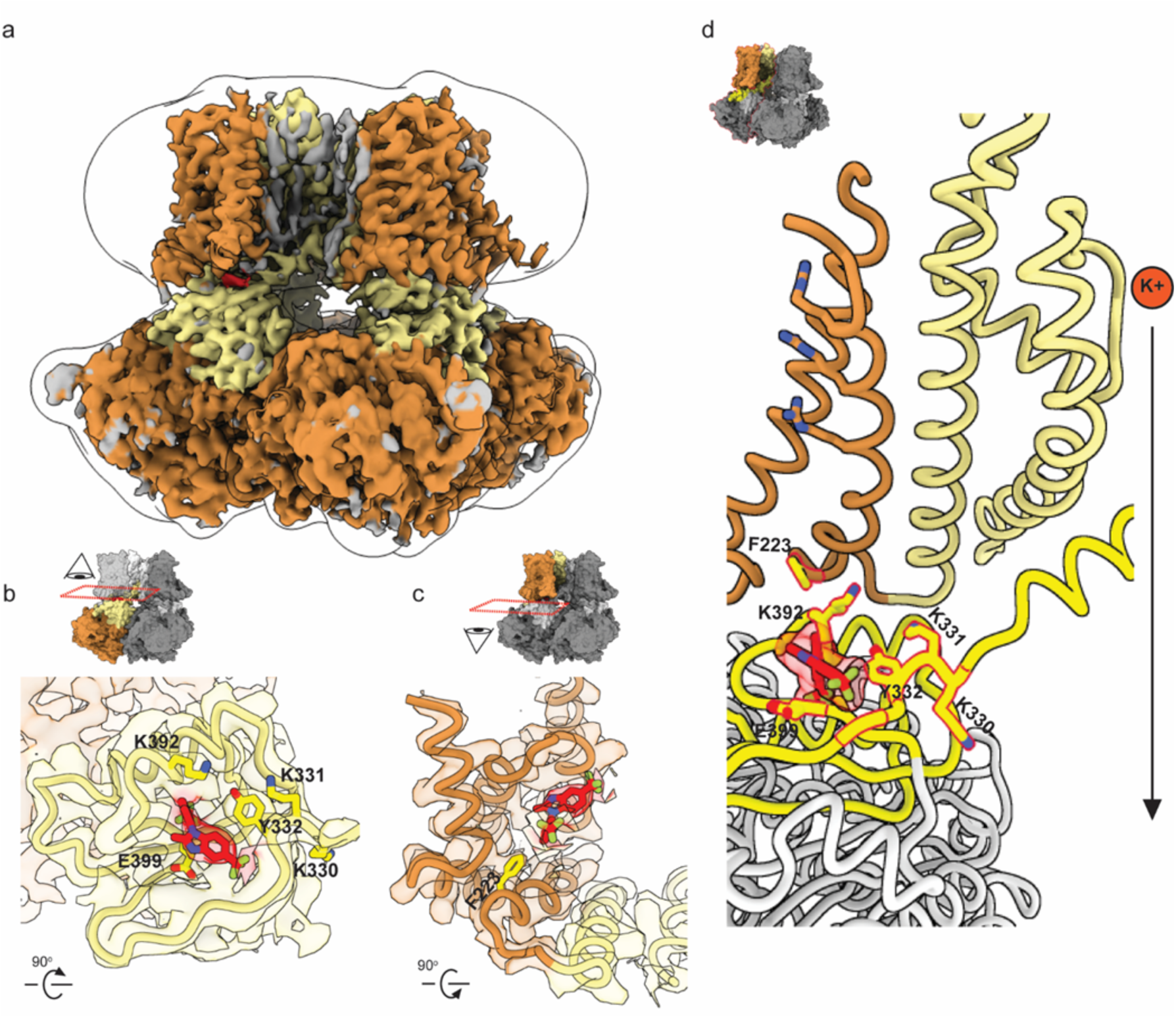
Coulombic Density of the hBK-NS1619 Complex. **(a)** This panel displays the density map of the BK channel as obtained by cryo-electron microscopy (cryo-EM). The channel was purified using detergent in the presence of calcium, with NS1619 added just before freezing. The density attributable to NS1619 is highlighted in red. The average resolution is 3.44 Å. **(b-d)** Top (b), bottom (c), and side (d) views of the NS1619 density, highlighting amino acids within a 5 Å distance (Y332, K392, and E399) and the amino acids whose mutation to alanine decreases the activation effect (F223, K330, and K331).

### Contribution of Binding Pocket Residues in NS1619 Coordination

To determine the role of each amino acid in the NS1619 binding site, we conducted alanine scanning mutagenesis on the residues appearing within 5 Å of NS1619 and those predicted most frequently from computational docking. Macroscopic currents were measured for the wild type (WT) and each mutant BK channel using symmetrical potassium solutions in the inside-out patch configuration. Perfusion with saturating concentrations of NS1619 (≤ 30 μM) induced a significant leftward shift in the I(V) curve by −36 ± 5 mV (N=8) in WT BK (Fig.2 c). Noteworthy differences from WT were observed only in mutations F223A, K330A, and K331A, with shifts of −20 ± 3 mV (N=5), −7 ± 2 mV (N=4), and −26 ± 2 mV (N=3) respectively (Fig. 2 d-f). Notably, K330 was identified in a previous study by Gessner et al. (2012) as critical to the mechanism of Cym04 binding, suggesting a shared binding site and putative activation mechanism with NS1619.

**Fig. 2.**
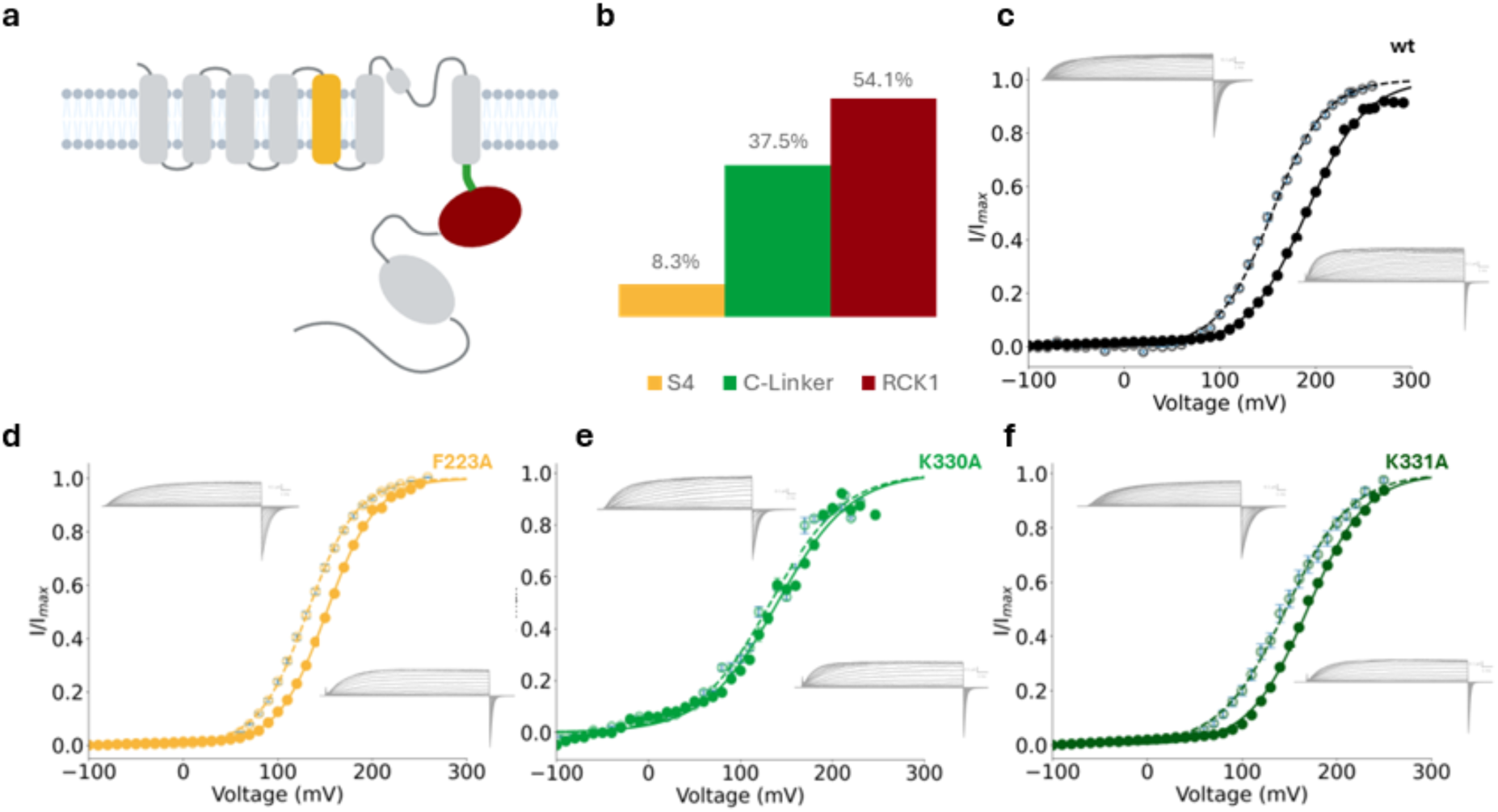
Characterization of the NS1619 binding site on the BK channel and effects of specific mutations. (a) Membrane topology representation of an α subunit of the BK channel, highlighting regions where molecular docking identified amino acids potentially involved in the NS1619 binding site. Regions are colored according to their involvement: voltage sensor domain (VSD) in golden yellow, C-linker in green, and RCK1 in maroon red. (b) Percentage distribution of 50 different docking poses for a total of 24 frequently repeated amino acids, where 8.3% belong to the VSD, 37.5% to the C-linker, and 54.1% to the RCK1. (c) Normalized current/voltage curves for the wild type (wt) BK channel in the absence (solid line) and presence (dashed line) of 30 µM NS1619, showing the effect of the activator on channel activity. (d-f) Alanine scanning analysis for amino acids that decrease the effect of NS1619, specifically for F223A (d), K330A (e), and K331A (f), demonstrating alterations in current responses in the presence of the activator.

We pursued computational docking protocols to identify the binding site of NS1619 on the BK channel based on our APO BK structure and the recently published cryo-EM structures of the hBK channel as templates (*23*). The docking evaluation of NS1619 was performed using a grid that includes the fully open (PDB: 6V38) and closed (PDB: 6V35) models in MAESTRO (Schrödinger Release 2022-2). NS1619 docking experiments revealed distinct patterns of optimal binding poses, identifying 24 implicated amino acids. Of these, 8.3% belong to the voltage sensor domain (VSD), 37.5% to the C-linker, and 54.1% to the RCK (Fig.2 a and b). We opted to test those amino acid residues whose frequency exceeded 50% in the pore domain. In the VSD we chose the amino acid with the highest occurrence (Fig. S3 a). Molecular docking also suggested a set of residues within the C-linker and RCK1 that potentially interact with NS1619, whose appearance frequency was equal to or greater than 50%, including R329, Y332, G334, F391, K392, and E399. However, mutating these residues to alanine resulted in a similar activation gating as that of WT-BK or subtle increases in the effect of NS1619, as shown in Figure S3 b-g.

### Dynamic Binding and Conformational Changes of the NS1619 Binding Pocket

To determine whether the binding site of NS1619 indeed undergoes a conformational reorientation, we conducted molecular dynamics simulations of the BK-NS1619 complex starting from the predicted closed channel configuration and running for 4.5 μs until reaching the open channel configuration in the presence of NS1619. In various simulations, NS1619 initially interacts with the residues identified from the BK-NS1619 complex structure (Fig. 3a, left panel). As the trajectory progresses, the S6 helix adopts an open conformation, allowing interaction of NS1619 with the K330-K331 and Y332 complex (Figure 3a, right panel). The side chains of both K330 and Y332 rotate following NS1619, while F315 in S6 adopts its open conformation. Both lysines (K330 and K331) interact with NS1619 through the aliphatic carbons of their side chains via van der Waals interactions, not with the amino groups. The activator finally interacts with F223, linking the S6 helix to S5, thereby stabilizing the open conformation of the channel, consistent with several molecular dynamic simulations.

**Fig. 3.**
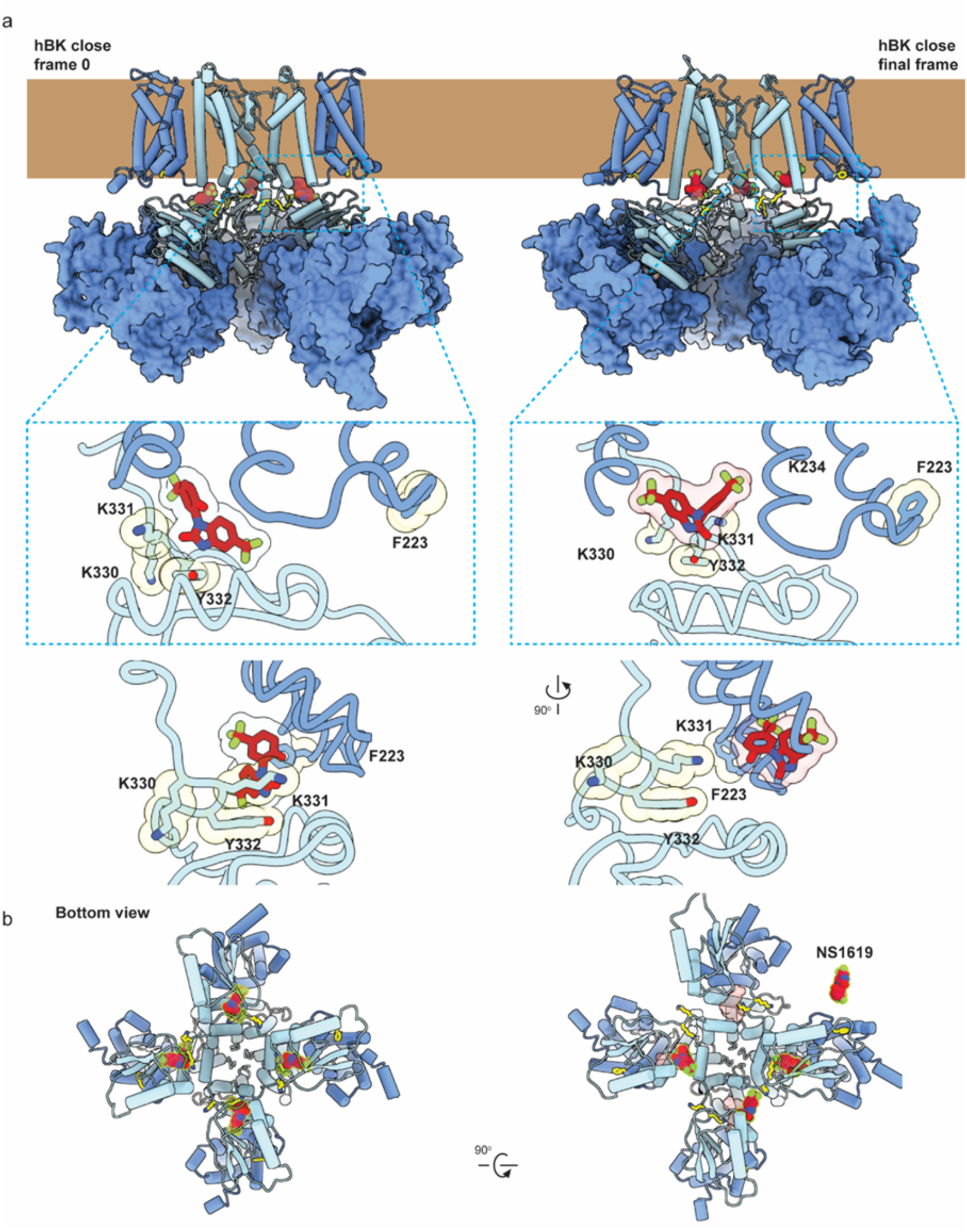
Molecular Dynamics Simulations Reveal the ‘Foot-in-the-Door’ Mechanism of NS1619 at the S6-RCK1 Interface of BK Channels. (a) Molecular simulation starts with the channel in a closed state (left panel) in the presence of NS1619 (highlighted in red), progressing to the open state (right panel). This sequence demonstrates how NS1619 facilitates the transition of the BK channel from its closed to open conformation. (b) Bottom view.

### NS1619 increases gate equilibrium constant Lo

Residues from boththe voltage sensor and pore domain interact with NS1619 as part of its binding pocket (Fig. 1,2). To quantify the involvement of pore gating in the activation mechanism of the BK channel by NS1619, we took advantage of the BK allosteric gating model (*24*), where the open probability (*P_o_*) in the absence of Ca^2+^ is given by the expression:

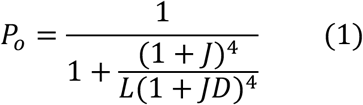

where, 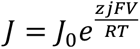 defines the equilibrium constant of the resting-active equilibrium of the voltage sensor, *J_0_* is the equilibrium constant at 0 mV, *z_J_* defines the voltage dependence of the voltage sensor activation, *V* is the applied voltage, and *F, R,* and *T* have their usual meanings. Eq. 1 becomes *Po = 1/(1 + 1/L)* when all the voltage sensors are resting, where 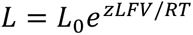. *L _o_* is the equilibrium constant defining the closed-open equilibrium at 0 mV, and *z_L_* is the voltage dependence for channel opening. Therefore, if the number of channels in the patch is known, *L* can be determined by plotting the absolute *Po* at sufficiently negative voltages (limiting slope method (*25*, *26*)). The number of channels in the patch can be obtained using non-stationary noise analysis. (Eq. 4) *z_L_* can be calculated in the absence of Ca^2+^ at very negative voltages using the relation

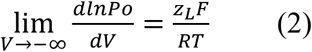

Figure 4a shows representative unitary currents at −60 mV without the activator, and (Figure 4b) in the presence of 30 μM NS1619, where the presence of the activator leads to an apparent increase in *Po*. *Po* data at negative voltages were plotted on a logarithmic scale (Figure 4f), and the slope of the straight lines fitting the data defines *z_L_*. Extrapolating this straight line helps determine *L_o_*, represented by the log Po data at zero mV (Fig. 4e-f). Under these conditions, *L_o_* is 4 × 10^-6^ ± 2 × 10^-7^ and 2 × 10^-5^ ± 1 × 10^-6^ in the absence and the presence of 30 μM NS1619, respectively.

**Fig. 4.**
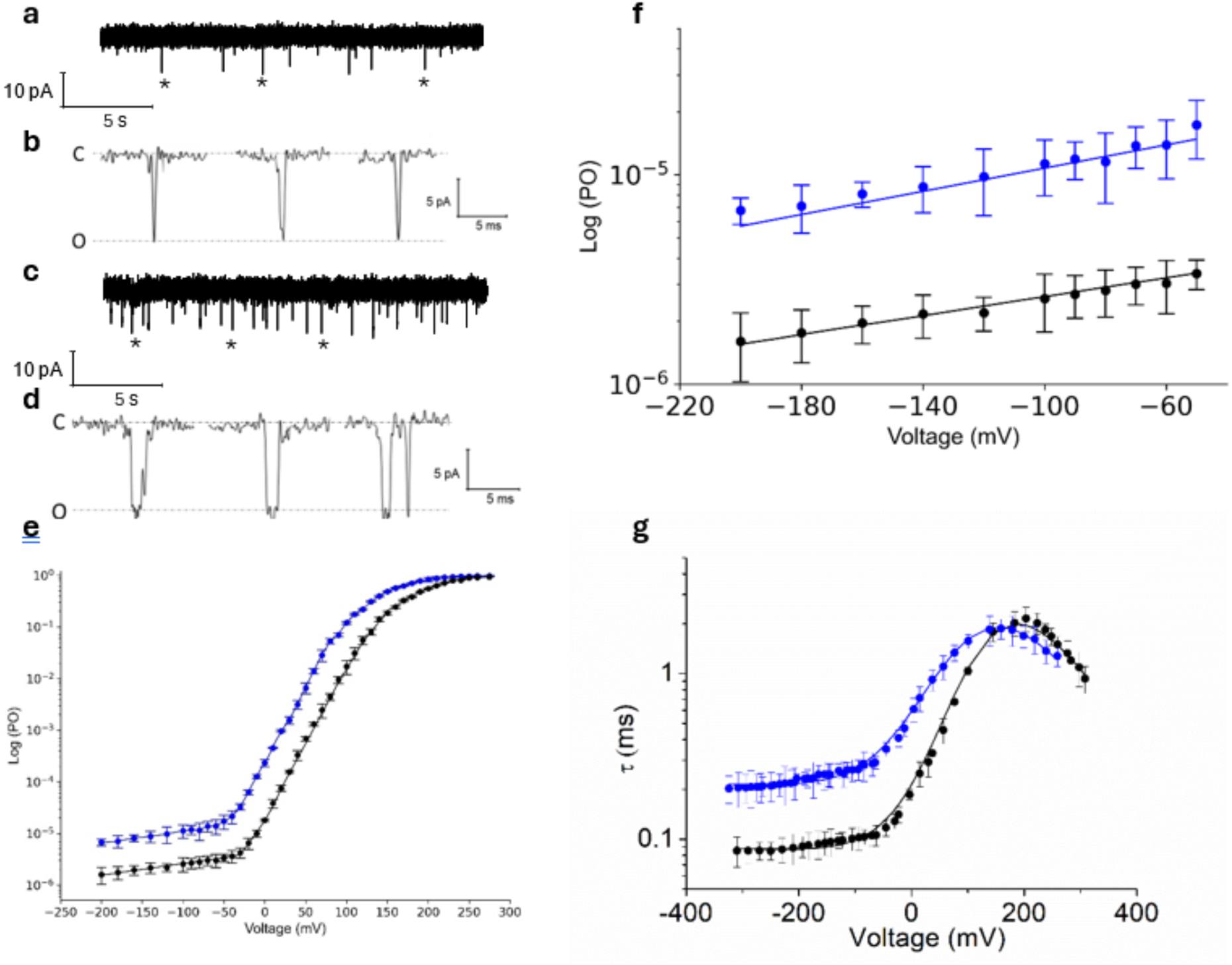
NS1619 Modulates the Equilibrium Constant L between the Open and Closed States of the BK Channel. **(a)** Representative BK currents in at −60 mV, with 0 µM NS1619. NPo = 6.018 × 10−4. **(c)** BK currents from the same patch as in a, following the addition of 30 µM NS1619. NPo = 5.092 × 10−3. The patch was estimated to have 489 active channels, determined by nonstationary noise analysis. **(b and d)** Representative channel openings from the traces in a and c, (from openings indicated by *, shown at an expanded time scale. **(e-f)** Logarithmic representation of Po vs. V for the BK channel without the activator (in black, N=6) and with 30 µM NS1619 (in blue, N=5). The solid lines represent the best fit using the HA model. Lo=4 × 10^-6^ ± 2 × 10^-7^ without the activator and 2 × 10^-5^ ± 1 × 10^-6^ with 30 µM NS1619. D=17 ± 2 and 8.5 ± 1 in the absence and presence of the activator, respectively. **(g)** τ versus voltage from patches with 0 (black circles) or 30 µM NS1619 (blue circles). Lines represent fits with Scheme 1 using parameters in Table 1.

Fig. 4g shows the time constant versus voltage in the absence (black circles) and presence (blue circles) of 30 µM NS1619, derived from macroscopic currents (Fig. S4). Notably, at negative potential, where the time constant is given by 1/γ, the time constant in the presence of the activator (τ) is higher, indicating that the rate describing the transition from the open to the closed state of the channel (γ), (Scheme 1) decreases. Since *L_0_ = δ_0_/ γ_0_*, where *δ_0_* is the forward rate constant, we conclude that all the change in *L* due to the agonist resides in the decrease in *γ*. To explore whether the change in γ alone is sufficient to explain the effect of NS1619 or if other parameters of the allosteric model (Scheme 1; (*27*)) are also involved, we fitted the logPo(V) data using equation 1. We found that the best fit (Fig. 4g, black and blue lines) is achieved by a decrease in both γ and the allosteric factor D (allosteric coupling between the PD and VSD), as shown in Table 1. Importantly, the rates describing the voltage sensor (α and β) remain unchanged in the presence of the activator. This aligns with the findings of Gessner et al (*22*), who demonstrated that the activator does not affect the channel’s Q/V relationship. Furthermore, the values obtained for *δo* and *γo* from τ are consistent with the Lo values derived from the limiting slope method, validating the hypothesis that NS1619 modulates this equilibrium constant.

These findings support our hypothesis that NS1619 activates BK by stabilizing the open conformation of the pore domain. Our data strongly support the idea that NS1619 influences the conformational stability of the PD, resulting in a longer channel open time and a decrease in the deactivation rate.

## Discussion

This study integrates techniques from electrophysiology, molecular docking, cryo-electron microscopy, and molecular dynamic simulations to define the binding site and mechanism of action of the BK channel activator NS1619. Single particle cryoEM and molecular dynamics analysis reveal that amino acids from the RCK1 and C-linker initially form the binding site. Upon NS1619 binding, the binding pocket reconfigures to involve an amino acid from S4 (VSD). This dynamic behavior allows NS1619 to stabilize the BK’s open conformation, highlighting a dual pathway of modulation that should help define new therapeutic targets.

Structural and molecular docking results point to a set of residues in the VSD, RCK1, and C-linker that putatively participate in NS1619 binding, including F223, R329, K330, K331, Y332, G334, F391, K392, and E399. However, side chain perturbations by mutagenesis indicate that only F223, K330, and K331 significantly alter the response to the activator compared to the wild type. This apparent discrepancy between structural data and functional results can be explained by considering the nature of these particular protein-ligand interactions. Although residues such as Y332, G334, F391, K392, H394, and E399 appear structurally close to NS1619, mutating these to alanine does not alter the channel function in the presence of the activator, suggesting their role may be more related to the structural conformation of the binding site or stabilization of the electrostatic milieu, rather than directly participating in the interaction with the activator. Our molecular dynamics results (Fig. 3) show that twisting the S6 segment induced by the final conformation of NS1619 in the binding pocket allows direct interaction between the K330-K331 complex and the activator, which also interacts with F223. These results suggest a “foot-in-the-door” mechanism where NS1619 would remain bound to F223 of the VSD on one side and to K330-K331 of the S6-RCK linker on the other, stabilizing the open configuration of the channel.

The reduction in NS1619 facilitation by these mutants correlates with the impact of the mutation on the closed-open pore equilibrium, defined by the equilibrium constant *Lo*. We note that K330A increases *Lo* approximately 24-fold, an effect on *Lo* much larger than the induced by mutations F223A and K331A, which increase *Lo* about 3-fold compared to the wild type (*28*). These differences suggest that K330 may play a crucial role anchoring NS1619 to the binding site and in transducing the conformational change that favors the opening of the channel. Mutant K330A likely alters the interaction dynamics between the transmembrane and cytosolic domains, preventing an efficient transition to the open conformation stabilized by NS1619. In contrast, although mutants F223A and K331A disrupt activation by NS1619, they lead to minor changes in *Lo*, indicating that their contribution to the stability of the open state is significant but less critical than that of K330. This analysis highlights how specific structural modifications within the BK channel can dramatically influence its pharmacology, underscoring the importance of these residues in the modulation of the channel by NS1619 and providing a direct link between the molecular structure of the channel and its biophysical function. A detailed understanding of these mechanisms is crucial for the biological characterization of the channel and in the structural-based design of more effective and specific modulators.

### A Common Binding Site in BK for Diverse Activators

The NS1619 binding site within the BK channel appears to act as a general druggable site for various activators, including NS004, NS1608, GoSlo-SR-5-6, CYM04, and GoSlo-SR-5-44. Molecular docking results consistently show that these compounds bind in the same binding pocket at NS1619, suggesting a shared and multifunctional site (Fig. S5). This finding is fundamental to understanding how various modulators can influence BK channel activity through a common mechanism and allows for designing new modulators that leverage this strategic interaction site. Molecular docking experiments demonstrate that the same amino acids implicated in the proposed binding site for NS1619 are involved in interactions with other activators like NS004, Cym04, NS1608, GoSlo-SR-5-44, and GoSlo-SR-5-6. These results underscore the versatility of the binding site to adapt to different molecular structures, emphasizing its capacity to interact with a wide range of compounds, both hydrophobic and polar. Our findings are consistent with previous studies, such as those by Webb et al. (*29*), who suggest that the binding site for GoSlo_SR-5-6 would be a hydrophobic pocket between the S4/S5 and S6 loop of the BK channel, reinforces the validity of our models and the therapeutic relevance of this approach.

### Activation Mechanism of the BK Channel by NS1619

Molecular dynamics, limiting slope, and time constant analysis all suggest that NS1619 activates the BK channel by shifting the closed-open equilibrium toward the open configuration, increasing the equilibrium constant *Lo,* and decreasing the allosteric factor *D*. It has been established that NS1619 does not affect the resting-active equilibrium of the voltage sensor and that the activation effect is independent of Ca^2+^ binding (*22*). Limiting slope fits shows an increase in the equilibrium constant between the closed and open states in the presence of the activator. When we evaluate the time constant at very negative potentials where τ is equivalent to 1/γ (describing the transition from open to closed), we observe a reduction in γ indicating that the channel closures slow down in the presence of NS1619 (Fig. 4f). Since the rate constant describing the transition from closed to open (δ) does not change in the presence of NS1619, the increase in *Lo* is entirely explained by the decrease in γ. Molecular dynamics calculations and our alanine scanning results suggest that the activator interacts with phenylalanine 223 in the VSD, an observation supported by the observed effect on the allosteric factor *D*.

Furthermore, molecular dynamics also show that once NS1619 binds to the final conformation of the binding site, a likely movement in S6 leads to an interaction with lysine 330. The fact that the K330A mutation eliminates the activating effect of NS1619 and the critical role of the interaction between NS1619 and K330 in stabilizing the channel in its open state leads us to propose a “foot-in-the-door” mechanism. In this scheme, the activator binds to its binding site. It undergoes a reconfiguration that enables NS1619 to interact with F223 of the VSD on one side and with the K330-K331 complex of the C-linker on the other, effectively stabilizing the open state of the channel and acting as a crucial ‘foot in the door’ (Figure 5). This interaction explains the observed effects on the equilibrium constant *Lo* and suggests that once the activator engages with K330, it significantly slows down the channel closing process. The present results are also aligned with the idea that other activators might share this binding site and mechanism, either partially or fully. For example, Webb et al. (*29*) found that GoSlo-RS-5-6 activates the channel by increasing *Lo, Jo*, and reducing *D*. Gessner suggests that Cym04 and NS1619 are modulators of *Lo* and *Jo*, shifting the equilibrium toward the open and active configuration respectively (*22*). However, Hoshi and Heinemann (*30*) argue that there is some ambiguity regarding *Jo*, as part of the effect could be due to changes in the coupling between the VSD and the ion conductance gate. As Gessner shows, the Q/V of the channel does not change in the presence of the activator, implying that the activator does not modify the voltage sensor resting-active equilibrium. Thus, the mechanism would be through an increase in *Lo* and a decrease in *D*, which aligns with our results. Furthermore, Rockman et al. propose that NS11021 activates the BK channel through the PD by slowing the rate constant between the open and closed states of the channel, concluding that NS11021 stabilizes an open state (*31*).

**Fig. 5.**
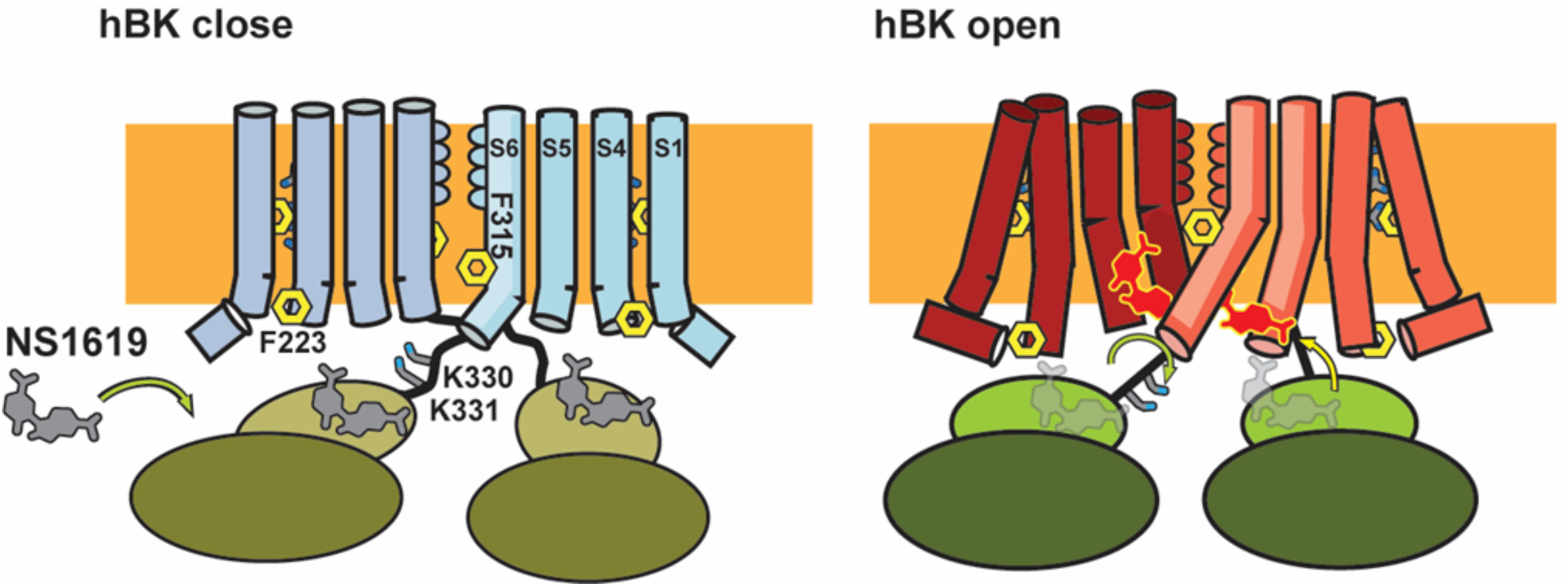
Mechanism of Activation by NS1619 in the BK Channel. This illustration depicts the proposed mechanism of activation of the BK channel by NS1619. Initially, NS1619 binds to its site on the channel. Following binding, a reconfiguration occurs, allowing NS1619 to simultaneously interact with F223 on the voltage sensor domain (VSD) and the K330-K331 complex on the C-linker. This interaction induces a positional shift of the S6 segment, effectively acting as a “foot-in-the-door” that stabilizes the channel in its open state.

## Conclusion

In this work, we have enhanced our understanding of the activation mechanism of the BK channel by the NS1619 activator, identifying a binding site whose active conformation stabilizes the open configuration of the channel, thereby preventing the transition to closure. This discovery sheds light on the regulation of ion channels and suggests a dual modulation pathway that could be therapeutically exploited. The involvement of specific residues in channel modulation, coupled with the proposed “foot-in-the-door” mechanism, highlights the complexity of ion channels as pharmacological targets and underscores the need for a thorough understanding of their modulation to design future drugs. Furthermore, the fact that this site may be common to various BK channel activators opens new avenues for developing innovative modulators that can leverage this strategic interaction site. A poorly selective hydrophobic binding site paves the way for the design of selective modulators, prioritizing hydrophobicity as a key factor contributing to the design of new drugs.

## Materials and Methods

### Channel Expression for electrophysiology

BK channels were expressed in *Xenopus laevis* oocytes. The cDNA for the wild-type human BK α-subunit (Genbank U11058) was kindly provided by L. Toro (University of California, Los Angeles, CA). Mutants were created using standard molecular biology techniques and verified by sequencing. mRNA was synthesized in vitro using the mMESSAGE mMACHINE kit from Ambion. Oocytes were then injected with 50 ng of mRNA and incubated in ND96 solution (NaCl, KCl, CaCl_2_, MgCl^2^, and HEPES, pH adjusted to 7.4 with NaOH) at 18°C for 3–7 days before electrophysiological analysis.

### Electrophysiological recordings

All current recordings were acquired using the patch-clamp technique in the inside-out configuration. Data was acquired with an Axopatch 200B (Molecular Devices) amplifier and the Clampe× 10 (Molecular Devices) acquisition software. The voltage command and current output were filtered at 10 kHz with an 8-pole Bessel low-pass filter (Frequency Devices). Current signals were sampled with a 16-bit A/D converter (Digidata 1550B; Molecular Devices), using a sampling rate of 500 kHz. Unless otherwise stated, linear membrane capacitance and leak subtraction were performed based on a P/-8 protocol (Bezanilla and Armstrong, 1977). Borosilicate capillary glasses (1B150F-4, World Precision Instruments) were pulled in a horizontal pipette puller (Sutter Instruments). After fire-polishing, pipette resistance was typically 1-2 MΩ. The ground electrode was an agar bridge containing 1M KMeSO_3_. All experiments were performed at room temperature (20-22 °C).

### Macroscopic currents

Potassium currents were elicited by pulse protocols with a holding potential of −100 mV followed by a test pulse to different voltage steps from −100 to 300 mV in 10 mV increments, separated by intervals of 10 ms, and ending with a voltage pulse of −100 mV. The pipette and bath solutions were the same and contained 110 mM KOH, 10 mM HEPES, 2 mM KCl, 1 mM EGTA (free Ca^2+^ ∼0.5 nM), and the pH was adjusted to 7.4 with methanosulfonic acid. All data analyses were conducted using Clampfit 11.0.3 (Molecular Devices), Python, and Excel 2013 (Microsoft). The voltage dependency of the macroscopic tail currents was quantified using a Boltzmann function.

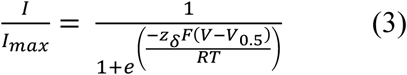

where *I_max_* is the maximum current, *V_0.5_* is the voltage at which *I/I_max_* equals 0.5, *F* is Faraday’s constant, *R* is the gas constant, *T* is the temperature in Kelvin, and *z_δ_* is the voltage dependency.

### Nonstationary Noise Analysis

A nonstationary noise analysis was applied, deriving the mean and variance 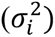 from isochronal segments of 200 current traces at 200 mV. We determined the unitary current (*i*) and the number of channels (*N*) by fitting the variance against the mean current using the expression

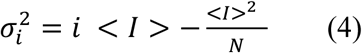

where <*I*> the mean current. Variance and mean current were determined using analysis software, which accounts for the gradual rundown of the current during experiments (*38*, *39*).

This method also allowed us to calculate the maximal *Po* (*Po max*) at 200 mV, using the maximum mean current obtained divided by *iN* (all channels open).

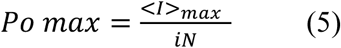

### Limiting Slope

Limiting slope experiments were performed using macropatches with hundreds of channels. *N*, *i* and *Po max* were determined as described in the previous section. After obtaining *N*, macroscopic currents induced by voltage pulses from −100 mV to 250 mV in increments of 10 mV were recorded. Voltage pulses were separated by 10 ms intervals. Unitary currents were recorded over 2–10 seconds, applying voltages ranging from −20 mV to −200 mV in steps of 20 mV. Both macroscopic and unitary current recording protocols were replicated on the same patch after perfusion with a solution containing 30 μM NS1619.

Histograms were created for all data points to obtain the NPo from single-channel current recordings. These histograms were normalized by dividing the ordinate values by the integral along the membrane current (*I*) axis to derive the probability density function (PDF), which delineates the likelihood of a given current level occurring based on the data distribution (*40*). Assuming that a maximum of nine channels could be open concurrently, the theoretical PDF was constructed from the sum of ten Gaussian distributions along the current axis.

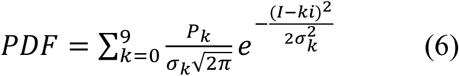

where *k* is the number of simultaneously open channels, and *i* is the unitary current. Each Gaussian distribution is centered around the expected current level, *ki*, with a variance of 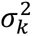. The variance for each current level accounts for both instrumental noise and the noise from the open channels. The overall variance 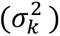 for the level with *k* open channels is taken as (*σ*_c_ + *kσ*_o_)^2^, where 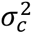 is the variance of the closed level, and 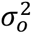 is the variance from the noise of an open channel. *P_k_* is the relative weight of each current level and was calculated using the expected probability of finding *k* open channels simultaneously on a membrane containing *N* channels, with an opening probability (*Po*), as described by a Poisson distribution (*40*).

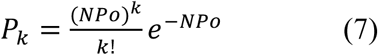

NPo was derived by fitting the theoretical PDF to the experimental PDF using the Solver add-in in MS-Excel. The adjustable parameters included *NPo*, 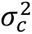, 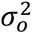, and *i*. The Po value was obtained by dividing by the number of channels determined from the nonstationary noise analysis conducted on the same patch.

### Activation and deactivation kinetics

The activation kinetic characteristics were obtained from macroscopic currents evoked in the inside-out configuration from −100 mV up to 300 mV in 10 mV steps of 10 ms duration with a prepulse of −100 mV and returning to −100 mV under symmetric potassium conditions. The protocol to assess deactivation consisted of a depolarizing pulse of 150 mV followed by test pulses from 0 to −300 mV in 10 mV steps and 5 ms duration. Both protocols were repeated after perfusing the internal side of the patch with a solution containing 30 µM NS1619. Current traces from activation and deactivation were quantified using a simple exponential from which the experimental time constant (τ) was derived. *τ* data was fitted using a 10-state voltage-dependent model (Scheme 1) that utilized microscopic rate constants instead of equilibrium constants, allowing for the calculation of activation and deactivation time constants for BK channel gating. This approach is analogous to the methods previously (*27*, *31*, *41*) using the equation:

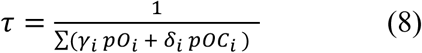

where *pO*_6_ and *pC*_6_ are the conditional occupancies of the open and closed states, respectively, *γ*_6_ is the rate describing the transition from the open to the closed state and *δ*_6_ is the rate describing the transition from the closed to the open state.

**Scheme 1.**
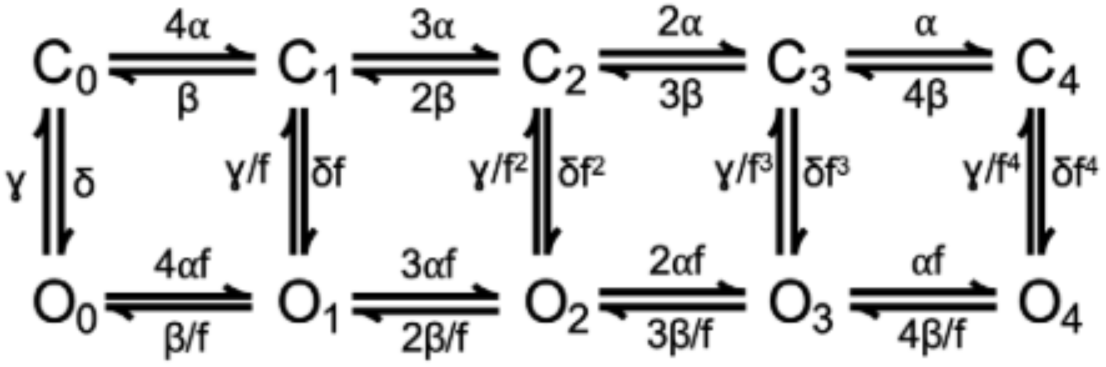

In Scheme 1, rate constants were defined by rates for VSD activation and deactivation (α and β, respectively), PD closing and opening (γ and δ), a gating valence for each of these rate constants (zα, zβ, zγ, and zδ), and allosteric coupling between the VSD and PD (D) using the additional simplifying assumption D = f^2^ (*31*).

### Molecular Docking

Using the electron microscopy structure of human BK (PDB: 6V35), we focused on the entire protein and employed the SiteMap module within the Schrödinger suite to identify potential ligand-binding sites. The BK protein was pre-processed and analyzed using SiteMap’s default settings to assess properties such as exposure, pocket volume, and hydrophobicity. This analysis generated contour maps (site maps), which we evaluated using Site Score values. For each identified site, we created receptor grids for docking, using cubic grids measuring 20 Å on each side. The most promising site, yielding the highest scores, was located near residues Ly331, Lys234, Arg326, and Ile322 on a single chain. We centered the grid on the centroid of each site map. Ligand docking was performed using the Glide program from the Schrödinger suite (2022–2) with default settings and the OPLS4 force field. The ligands were initially sketched in 2D and prepared using the LigPrep suite. We conducted the docking procedures in standard precision (SP) mode, which allows for flexible ligand sampling. The poses were scored and sorted using the default Glide Docking Score function, and the configurations with the most favorable docking scores were selected, confirming this site as druggable.

| compound | Score Kcal/mol |
| --- | --- |
| NS1608 | -5.3 |
| NS004 | -5.0 |
| SR-5-44 | -4.8 |
| CyM04 | -3.2 |
| NS11021 | -3.1 |
| SR-5-6 | -4.6 |
| NS1619 (6V22, open) | - 6.1 |
| NS1619 (6V35, closed) | - 4.4 |

### Molecular Dynamics Simulations

We used the 3.5 Å resolution Cryo-EM structures of BK channels, with entry code 6V35 representing the closed state and 6V22 representing the open state (*23*). To complete and refine small loops within these structures, we employed AlphaFold (*42*). The proteins were integrated into a POPC (1-palmitoyl-2-oleoyl-phosphatidylcholine) membrane system, which consisted of 785 lipids, 125,611 water molecules, and ions (338 K+, 398 Cl−), with a salt concentration of 0.15 M. The final system dimensions were 180 x 180 x 181 Å³. Molecular dynamics (MD) simulations were conducted using the AMBER software package (*43*). The protein structures were initially constrained in the loops with a harmonic restraint of 10 kcal·mol⁻¹Å⁻². The system underwent a relaxation process, where the restraints were progressively reduced by applying decreasing force constants in the following order: 10, 5, 2.5, 1.0, 0.5, and finally 0 kcal·mol⁻¹ Å⁻². This stepwise reduction facilitated a smooth transition to an unrestrained state, ensuring the stability and equilibrium of the system. 4.5 μs of MD simulations were performed with a 2 fs timestep for both proteins. Van der Waals interactions were calculated using a 1.0 nm cutoff, and a dispersion correction was applied for both energy and pressure (*44*). The particle mesh Ewald (PME) method was utilized for electrostatic interactions (*45*). The simulations employed the Amber19SB force field for proteins (*46*), the LIPID 21 force field for lipids (*47*), the OPC water model (*44*), and the ion parameters described previously (*48*). Temperature was maintained at 310 K using the velocity rescale (v-rescale) thermostat (*49*), while the semi-isotropic Berendsen barostat was employed to keep the pressure at 1 atm and 1 bar (*50*). Visual Molecular Dynamics (VMD) software carried out system construction and image generation (*51*). Histograms related to angles, distances, and displacements from the MD simulations were plotted using the Qtiplot program.

### Cloning, Expression and Purification for Cryo-EM experiment

A human Slo1 channel was cloned into the pFastbac vector containing a C-terminal 3C protease site, eGFP. P0 baculovirus was generated using the Bac-to-Bac method (Invitrogen) using Cellfectin II as the transfection reagent (Thermo Fisher Scientific, 10362100).

The P0 virus was amplified once to yield the P1 baculovirus, which was used to infect sf9 cells at a 5% (v/v) ratio. The cells were cultured at 27 °C in sf-900 II SFM medium (Invitrogen) for 48 h before harvesting. Cells were pelleted, washed with PBS, rapidly frozen in liquid nitrogen, and stored at −80°C until use. For purification, all steps were performed at 4°C and frozen cell pellets were thawed, diluted, and detergent-extracted in 20 mM Tris-HCl pH 8, 320 mM KCl, 10 mM MgCl_2_, 10mM CaCl_2_, 2 mM TCEP, 1% LMNG for 90 min. The solubilized supernatant was isolated by ultracentrifugation and incubated for 2 h with 1 ml CNBR-activated Sepharose beads (GE Healthcare) coupled with 4 mg high-affinity GFP nanobodies (*52*). Beads were collected by low-speed centrifugation, washed in batch mode with 10 column volumes of main buffer 20 mM Tris-HCl pH 8, 320 mM KCl, 10 mM MgCl_2_, 10 mM CaCl_2_, 2 mM TCEP, 1 μL/mL aprotinin, 1 μL/mL Pepstatin, 10 μL/mL soy tripsin y 100 μL/10mL 1-palmitoyl-2-oleoyl-glycero-3-phosphocholine (POPC):1-palmitoyl-2-oleoyl-sn-glycero-3-phosphoethanolamine (POPE): 1-palmitoyl-2-oleoyl-sn-glycero-3-phosphate (POPA) 5:5:1(PC:PE:PA) diluted in GDM, collected on a column by gravity, and washed with another 10 column volumes. Proteins were cleaved by the HRV 3C protease (*53*) overnight, concentrated, and analyzed using SEC on a Superose 6, 10/300 GE column (GE Healthcare), with running buffer containing 20 mM Tris-HCl pH 8, 320 mM KCl, 10 mM MgCl_2_, 10mM CaCl_2_, 2 mM TCEP, 1 μL/mL aprotinin, 1 μL/mL Pepstatin, 10 μL/mL soy tripsin y 50 μL/10mL PC:PE:PA. Peak fractions were collected and concentrated using a 100 kDa molecular mass cutoff centrifugal filter (Millipore concentrator unit) to 3.5 mg/ml. The concentrated protein was immediately used for the cryo-EM grid-freezing step.

### Cryo-EM sample preparation and imaging

Quantifoil 200-mesh 1.2/1.3 grids (Quantifoil) were plasma-cleaned for 30 s in an air mixture in a Solarus Plasma Cleaner (Gatan). Purified hBK-NS1619 samples were applied onto the grids and frozen in liquid-nitrogen-cooled liquid ethane using a Vitrobot Mark IV (FEI) and the following parameters: sample volume 3.5 μl, blot time 3.5 sec, blot force 3, humidity 100%, temperature 22 °C and double filter papers on each side of the vitrobot. Grids were imaged on a Titan Krios with a K3 detector (in super-resolution mode) and GIF energy filter (set to 20 eV) at a nominal magnification of ×130,000, corresponding to a super-resolution pixel size of 0.5315 Å, 0.55 Å or 0.56 Å per pixel depending on the default set-up at EM facilities mentioned above, respectively. The videos were acquired at 1 e−/ A^2^ per frame for 50 frames.

### Single-particle cryo-EM analysis

Image processing and three-dimensional map reconstruction were performed using the cryoSPARC program. Initially, individual particles were selected from the cryo-EM images and aligned and classified into 2D classes. Subsequently, 3D classification and refinement were carried out to reconstruct the Coulomb density map. Finally, a non-uniform refinement was performed applying C1 and C4 symmetry, achieving a final resolution of 3.83Å and 3.44 Å, respectively. Figure S2 shows a diagram of the overall processing and refinement strategy.

### Model building

The model of hSlo1 (PDB 6v38) was used to build atomic models for the hSlo1-NS1619 complex into our density maps. The initial models were built via iterative rounds of manual model building on COOT (*54*) and real space refinement in Phenix (*55*). The final refined atomic model was obtained using interactive, flexible fitting using ISOLDE (*56*). All structural analyses and figures were generated using UCSF ChimeraX (*57*).

## Acknowledgments

We thank Luisa Soto and Margaret Milewski for their technical assistance with molecular biology procedures. We are also grateful to Patrick Haller and Dr. Navid Bavi for their valuable help and guidance during the process of obtaining the Coulomb density maps.

## Author contributions

Conceptualization: RL, EP, NG-S, GFC

Methodology: NG-S, GFC, MR, YD, FDG-N, EP

Investigation: All authors

Visualization: All authors

Supervision: RL, EP, FDG-N

Writing—original draft: NG-S, GFC, RL

Writing—review & editing: NG-S, GFC, RL, EP

## Data and materials availability

All data are available in the main text or the supplementary materials. Cryo-EM maps and atomic models will be deposited in the Electron Microscopy Data Bank and the Protein Data Bank upon acceptance of the manuscript.

