## Supplemental Data 1 for "The BK channel-NS1619 agonist complex reveals molecular insights on allosteric activation gating"

Naileth Gonzalez-Sanabria *et al.*

; or.

**This PDF file includes:**

Figs. S1 to S5
Tables S1

Fig. S1

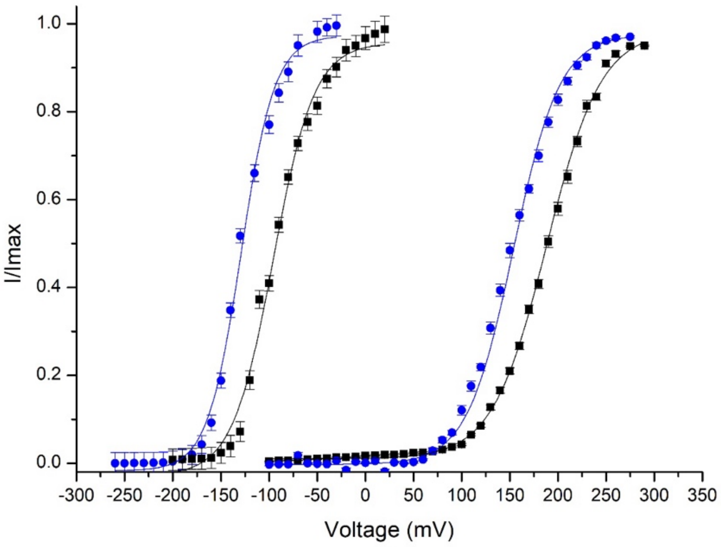


**Fig. S1. Effect of NS1619 in the absence and presence of 100 µM $CaCl_2$**. I/Imax vs voltage curves of the wild-type channel without activator or calcium in black solid line ($V_{0.5}$= 190 ± 6, $z_\delta$ = 0.90 ± 0.01). The continuous blue line corresponds to the I/Imax vs voltage curve in the presence of 30 µM NS1619 without calcium ($V_{0.5}$= 154 ± 7, $z_\delta$ = 0.91 ± 0.02). The black dashed line is in the presence of 100 µM $CaCl_2$, but without activator ($V_{0.5}$= -92 ± 4, $z_\delta$ = 1.03 ± 0.06), and the blue dashed line is in the presence of NS1619 and $CaCl_2$ ($V_{0.5}$= -129 ± 5, $z_\delta$ = 0.99 ± 0.02).

Fig. S2.

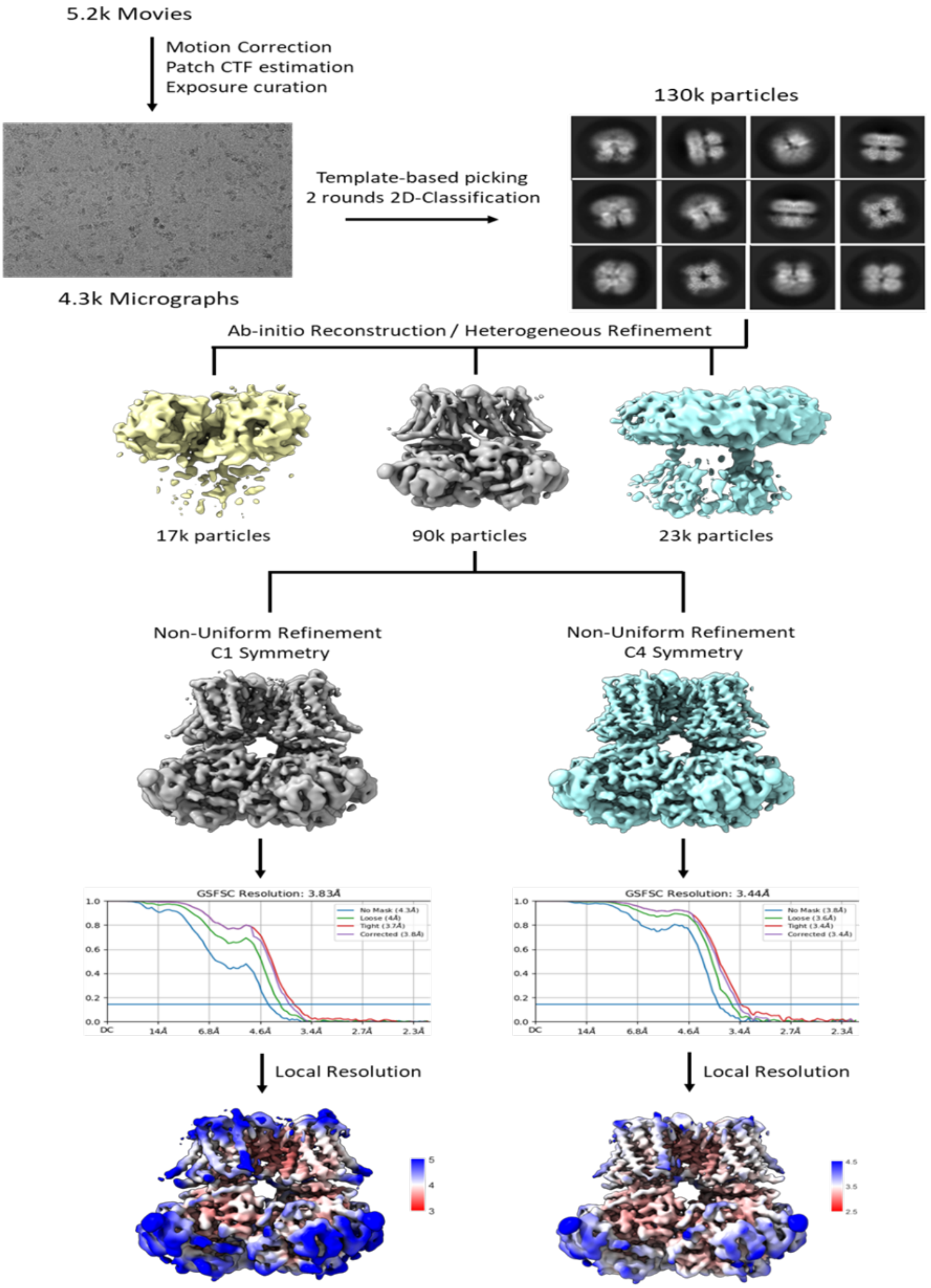


**Fig. S2. Diagram of global processing and refinement strategy.**

Fig. S3.

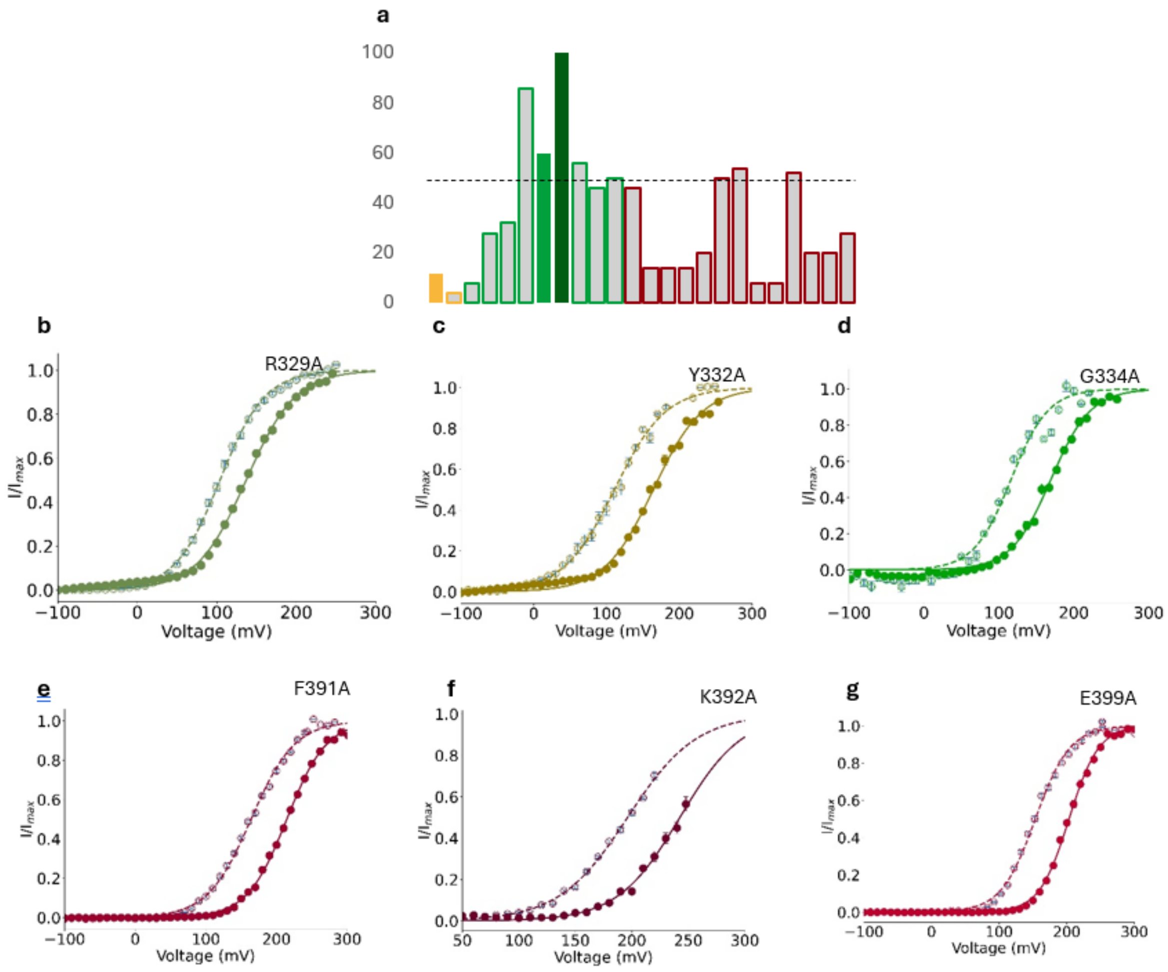


**Fig. S3. NS1619 docking frequencies and alanine scanning results. (a)** Frequency of occurrence of each amino acid in 50 dockings, from left to right: F223, N231, I326, G327, N328, R329, K330, K331, Y332, G333, G334, S382, P383, N384, L385, E388, F391, K392 , F395, V398, E399, F400, Y401 and Q402 the colors represent the area of the channel where each amino acid belongs: voltage sensor domain (VSD) in golden yellow, C-linker in green, and RCK1 in maroon red. The filled bars are the amino acids where the effect of NS1619 decreased in alanine scanning. The dotted line highlights 50%, only the amino acids that were equal to or greater than this value was evaluated **(b-g)** Alanine scanning analysis for R329A (b), Y332A (c), G334A (d), F391A (e), K392A (f) and E399A (g), note that in neither case was there a decrease in the effect of NS1619.

Fig. S4.

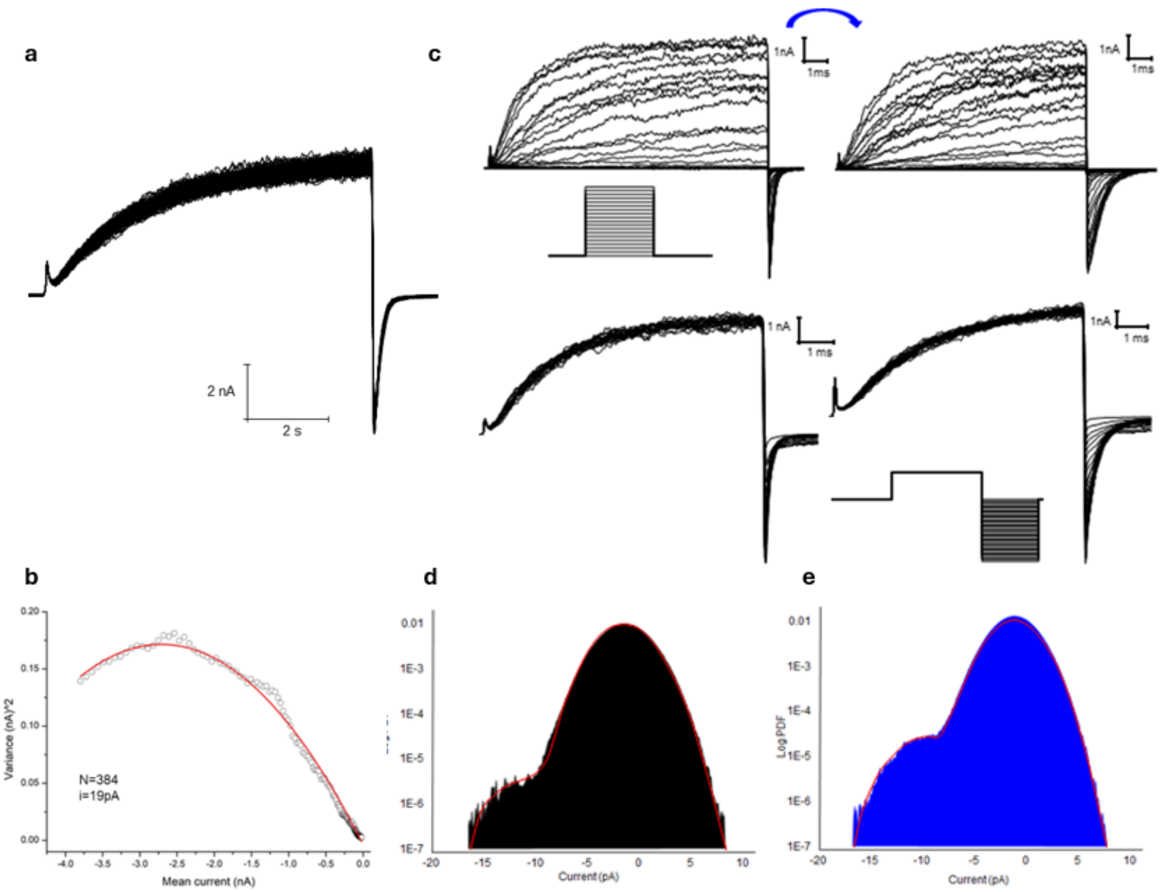


**Fig. S4. (a)** 200 macroscopic current traces evoked at 200 mV with a -100-mV tail and a holding potential of -100 mV. **(b)** Variance vs. mean current plot, fitted to Equation 4, with results of N = 384 and i = 19 pA. **(c, top panel)** Macroscopic current of BKwt in inside-out configuration, holding at -100 mV. The current was evoked by voltage pulses ranging from -100 mV to 300 mV in 10 mV increments, with a prepulse at -100 mV and returning to -100 mV, under symmetrical 110 mM potassium conditions. The current records on the left were obtained in the absence of NS1619, and those on the right in the presence of the agonist to a final concentration of 30μM **(c, bottom panel)** Deactivation protocol, holding at -100 mV. A depolarizing pulse of 150 mV was applied, followed by test pulses ranging from 0 to -300 mV in 10 mV steps. **(d-e)** Normalized histograms of all data points from the corresponding traces in Fig. 4a and 4b are shown (black for control, blue for NS1619). Red lines indicate fits to Equation 6, from which the NPo values for each voltage were extracted.

Fig. S5.

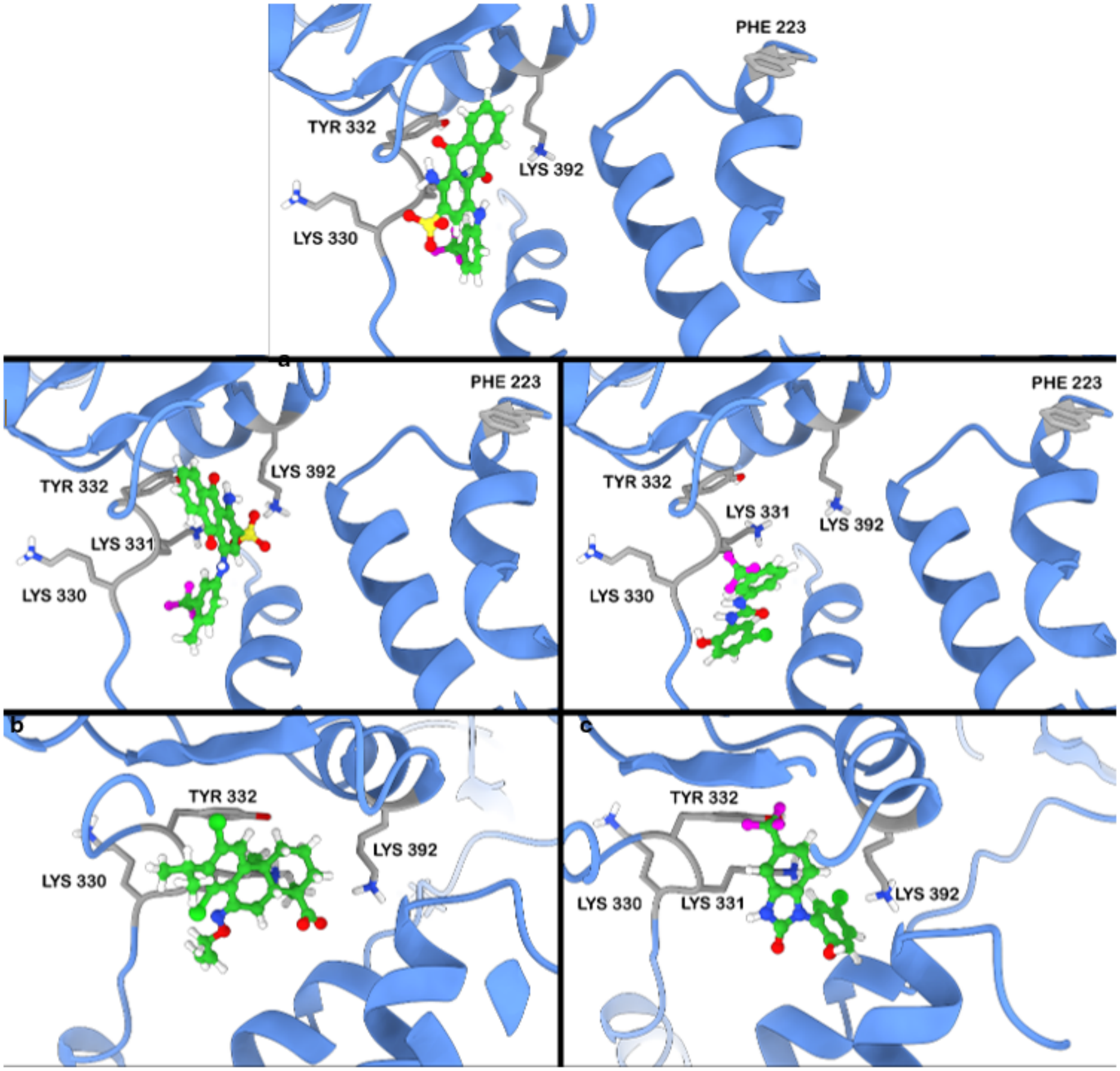


**Fig. S5. The Common Binding Site in the BK Channel for Diverse Activators.** Molecular docking using the PDB structure: 6V35 of the BK channel with different channel activators: **(a)** SR-5-6 **(b)** SR-5-44 **(c)** NS1618 **(d)** CyM04 **(e)** NS004.

**Table S1. Fitted parameters for Scheme 1 constrained by time constants acquired with and either 0 or 30 µM NS1619 at 0 $Ca^{+2}$**

| **Parameter** | **0 µM** | **30 µM NS1619** |
|---|---|---|
| α ($s^{-1}$) | 1608 | 1608 |
| β ($s^{-1}$) | 42744 | 42744 |
| $z_α$ ($e_0$) | 0.31 | 0.31 |
| $z_β$ ($e_0$) | -0.31 | -0.31 |
| δ ($s^{-1}$) | 0.03107 | 0.03152 |
| ϒ ($s^{-1}$) | 5379 | 1639 |
| $z_δ$ ($e_0$) | 0.32 | 0.32 |
| $z_ϒ$ ($e_0$) | -0.025 | -0.025 |
| D | 19.8 | 9.6 |
